# Targeting the *Plasmodium falciparum* UCHL3 ubiquitin hydrolase using chemically constrained peptides

**DOI:** 10.1101/2024.01.11.575158

**Authors:** Harry R. King, Mark Bycroft, Thanh-Binh Nguyen, Geoff Kelly, Alexander A. Vinogradov, Pamela J. E. Rowling, Katherine Stott, David B. Ascher, Hiroaki Suga, Laura S. Itzhaki, Katerina Artavanis-Tsakonas

## Abstract

The ubiquitin-proteasome system is essential to all eukaryotes and has been shown to be critical to parasite survival as well, including *Plasmodium falciparum*, the causative agent of the deadliest form of malarial disease. Despite the central role of the ubiquitin-proteasome pathway to parasite viability across its entire life-cycle, specific inhibitors targeting the individual enzymes mediating ubiquitin attachment and removal do not currently exist. The ability to disrupt *P. falciparum* growth at multiple developmental stages is particularly attractive as this could potentially prevent both disease pathology, caused by asexually dividing parasites, as well as transmission which is mediated by sexually differentiated parasites. The deubiquitinating enzyme PfUCHL3 is an essential protein, transcribed across both human and mosquito developmental stages. PfUCHL3 is considered hard to drug by conventional methods given the high level of homology of its active site to human UCHL3 as well as to other UCH domain enzymes. Here, we apply the RaPID mRNA display technology and identify constrained peptides capable of binding to PfUCHL3 with nanomolar affinities. The two lead peptides were found to selectively inhibit the deubiquitinase activity of PfUCHL3 versus HsUCHL3. NMR spectroscopy revealed that the peptides do not act by binding to the active site but instead block binding of the ubiquitin substrate. We demonstrate that this approach can be used to target essential protein-protein interactions within the *Plasmodium* ubiquitin pathway, enabling the application of chemically constrained peptides as a novel class of anti-malarial therapeutics.

## Introduction

Drug resistance to frontline malaria therapeutics is rapidly increasing, threatening our ability to limit the spread of the parasite and the debilitating disease it causes (1–5). As such, novel approaches to control parasite growth and transmission are urgently needed. The ubiquitin-proteasome system is essential to all eukaryotes and has been shown to be critical to *Plasmodium* survival across its entire lifecycle. Despite this central role to viability, specific inhibitors targeting the individual enzymes that mediate ubiquitin attachment (E1, E2, E3) and removal (deubiquitinases (DUBs)) do not currently exist. The ability to disrupt *P. falciparum* growth at multiple developmental stages is particularly attractive as this could potentially prevent both disease pathology, caused by asexually dividing parasites, as well as transmission, which is mediated by sexually differentiated parasites (6–8). The deubiquitinating enzyme PfUCHL3 is one such example. It is an essential protein, transcribed across both human and mosquito developmental stages. We have previously characterized this enzyme as a dual DUB/deNeddylase and determined its crystal structure in both Ub-bound and unbound states (9–12). PfUCHL3 is considered to be an ‘undruggable’ target given the high level of homology of its active site to human UCHL3 as well as to other UCH domain enzymes. As such, small molecule approaches to inhibit its activity would likely be cross-reactive. Moreover, the rest of the protein structure lacks defined pockets where small molecules could bind.

Small molecules and peptides aimed at disrupting protein-protein interactions have shown significant promise as therapeutics in both chronic and communicable diseases (13–17). An advantage of peptide-based drugs is the larger surface of interaction with their therapeutic target as compared to small molecules, which in turn leads to very high-affinity and specific binding and, thus, fewer off-target effects. Moreover, small molecules commonly require deep binding pockets in order to function as specific inhibitors (18) Peptides can target what is known as ‘undruggable’ proteins, those that lack a suitable binding pocket but have much larger and flatter surfaces where protein or nucleic acid partners bind (19–22). Short linear peptides (< 50 amino acids) containing only canonical amino acids make poor therapeutic compounds primarily due to their low biostability and bioavailability. Peptide macrocyclization and inclusion of non-canonical amino acids have been shown to substantially increase proteolytic stability, cell permeability, and binding affinity, and greatly expands the chemical diversity (21, 23, 24). Efforts to develop and deliver these types of drugs in the context of malaria infection are ongoing. There are several examples of peptides that have been found to inhibit *P. falciparum* proteins, such as the 13-residue β-hairpin peptide CFTTRMSPPQQIC that inhibits the interaction between the parasite’s Apical Membrane Antigen 1 (AMA1) and the host cell’s rhoptry neck protein (RON) thus blocking the parasite’s ability to invade new erythrocytes (25). Cyclomarin A is a cyclic heptamer found to inhibit *P. falciparum* diadenosine triphosphate hydrolase which inhibits parasite growth (26). Mahafacyclin B is a naturally occurring, cyclic heptamer identified from the latex of *Jatropha mahafalensis* and found to affect parasite viability during the asexual stage (27). A more recent example is a modified tetrapeptide, ‘3j’, containing a boronic acid warhead in place of the C-terminal carboxyl group which binds to the *P. falciparum* enzyme SUB1. The warhead attacks the active site Ser606 of PfSUB1 and covalently inactivates the enzyme which renders *Plasmodium* asexual-stage parasites unable to undergo egress, effectively blocking reinfection to other red blood cells (28).

Here, we apply the Random nonstandard Peptides Integrated Discovery (RaPID) display technology to identify constrained peptides capable of binding to PfUCHL3 with nanomolar affinities (16, 29–32). The two lead peptides were found to selectively inhibit the deubiquitinase activity of PfUCHL3 versus that of HsUCHL3. NMR spectroscopy revealed that the peptides do not act by binding to the active site but instead block the interaction of PfUCHL3 with the ubiquitin substrate. We demonstrate that these approaches can be used to target essential protein-protein interactions within the *Plasmodium* ubiquitin pathway, enabling the use of chemically constrained peptides as a novel class of anti-malarial therapeutics.

## Results

### mRNA display screening identifies peptides that specifically bind to PfUCHL3

In order to maximize the size of the peptide library, we used the RaPID platform, an in vitro mRNA display selection technique that circumvents the need for *in vivo* transformation steps and allows for the incorporation of nonproteinogenic amino acids to increase chemical diversity. During in vitro translation, each peptide is covalently linked to its cognate mRNA via a puromycin linker. After ribosomal release, the peptide-mRNA is reverse-transcribed to generate a peptide-mRNA-DNA complex. When scaled up and performed in a combinatorial format, libraries in the order of 10^14^ unique peptides can be generated and screened for biological activities of interest (**Figure 1**). By using ‘flexizymes’, small (45-46 nucleotides) artificial aaRS-like ribozymes, tRNA substrates can be aminoacylated with unnatural amino acids (UAAs) of choice (29). To conduct genetic code reprogramming in the RaPID system, the natural tRNAs and amino acids for the codon being reprogrammed are omitted from the translation mixture and replaced with engineered tRNAs for that codon which have the desired UAA preloaded onto them by the appropriate flexizyme. In our case, the mRNA library is designed to reassign N-formyl-methionine as N-chloroacetyl-D-tryptophan as the translation initiator. The peptides also feature a carboxy-terminal cysteine such that after translation, the peptides spontaneously cyclise forming a non-reducible thioether bridge (33). Macrocyclic peptides are more conformationally constrained compared to their linear counterparts, and therefore often possess higher substrate specificity and affinity. The non-reducible, D-stereochemistry cyclisation used in the RaPID system leads to the peptides with increased proteolytic stabilities. During RaPID, peptides undergo repeated rounds of selection, whereby they are screened against the protein of interest, in our case PfUCHL3, which had been immobilised onto streptavidin-coated Dynabeads via a biotin tag (**Supplementary Figure 1**). PfUCHL3 was cloned with an N-terminal AviTag^TM^, GLNDIFEAQKIEWHE, between the protein and a 6xHIS tag. Purified BirA enzyme recognizes the AviTag^TM^ sequence and was used to catalyse the monovalent biotinylation of PfUCHL3 at a location distal to the protein’s surface.

**Figure 1.**
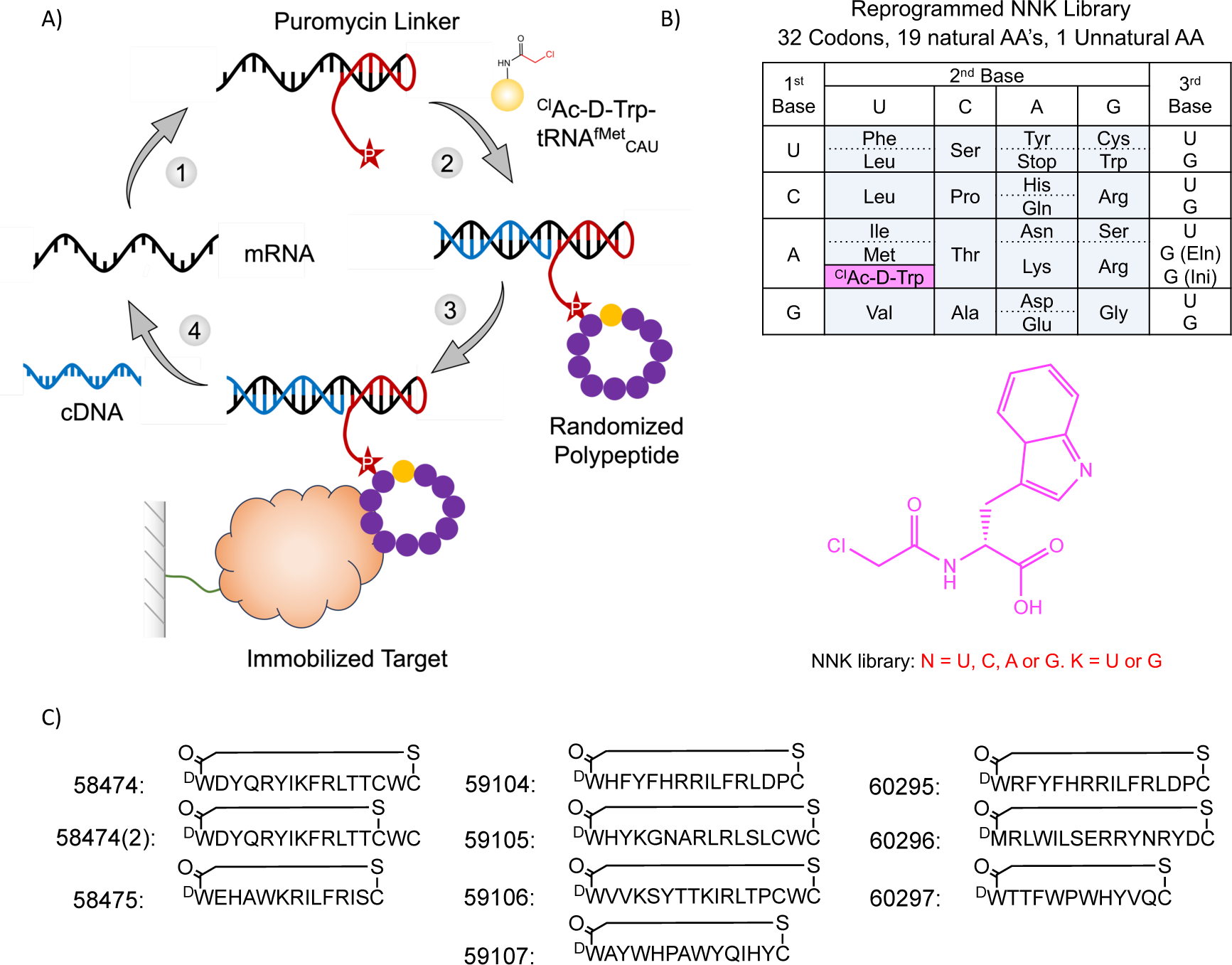
RaPID methodology peptide selection. A) In the RaPID technology, an mRNA library is ligated to a puromycin linker (1), translated and reverse transcribed (2), selected against an immobilised protein of interest (3), and cDNA recovered to sequence and regenerate a new mRNA library for a subsequent round of selection (4). B) Summary of the NNK genetic code used in the randomised region of the peptide library. The pink box shows the unnatural amino acid which was genetically reprogrammed to replace the initiator methionine (AUG (Ini)) using flexizyme technology, the elongator methionine (AUG(Eln)) was left unprogrammed. C) The top ten most enriched peptide motifs from selections against PfUCHL3.

Following seven rounds of selection against PfUCHL3, the peptide library had reached convergence, and ten peptides from unique peptide families that displayed high sequence enrichment as measured by the next generation sequencing were selected for synthesis (**Figure 1C**). The peptides ranged from 12-17 amino acids in length and were synthesized commercially in their cyclised state. Since peptides 59105, 59106 and 58474 contained an additional internal cysteine residue, there was potential for two cyclisation conformations. Peptides 59105 and 59106 appeared to purify solely as a single species, suggesting that one isomer is the more favourable (likely the peptide cyclising to the cysteine nearest the N-terminus) or that the two different cyclisation states co-elute and can not easily be separated. However, for peptide 58474 the HPLC trace revealed that both cyclisation states were present, and could be purified as two different peptides for downstream analysis, peptide 58474 (the major product) and peptide 58474(2) (the minor product).

### Macrocyclic peptides bind to PfUCHL3 with high affinity

The affinities and kinetics of binding of PfUCHL3 to the peptides were analysed by biolayer interferometry (BLI). Most of the peptides showed binding, albeit with a wide range of *K*_d_ values from low nM to mid-μM **(Table 1)**. Some peptides (59105, 59106, 60295 and 60297) displayed clear biphasic binding profiles that were fit to a heterogeneous binding model, while others (59484, 59104 and 60296) could be fit to a simpler 1:1 model. Inspection of the tightest *K*_d_ for each peptide indicated that 60296 and 60297 have the highest affinity (9 nM and 2 nM, respectively), followed by 58484 and 59105 (high nM), and the rest are in the μM range.

**Table 1.** Dissociation constants, on-rates and off-rates measured by BLI and thermal shifts measured by TSA.

| Peptide | $K_d$ | $k_{on}$ ($\times 10^4$ M $s^{-1}$) | $k_{off}$ ($\times 10^{-4}$ $s^{-1}$) | $\Delta T_m$ ($^{\circ}C$) PfUCL3<br>(+ 6 $\mu M$ peptide) |
| --- | --- | --- | --- | --- |
| 58474 | $101 \pm 7$ nM | $1.71 \pm 0.09$ | $17.4 \pm 0.9$ | $+1.2 \pm 0.14$ |
| 58474(2) | low | low | low | $+1.3 \pm 0.08$ |
| 58475 | low | low | low | $+0.5 \pm 0.15$ |
| 59104 | $170 \pm 5$ nM | $1.81 \pm 0.05$ | $30.7 \pm 0.4$ | $-0.4 \pm 0.04$ |
| 59105 | $\sim 9$ $\mu M^*$ | | | $+0.1 \pm 0.06$ |
| 59106 | $\sim 28$ $\mu M^*$ | | | $-0.4 \pm 0.03$ |
| 59107 | NSB | NSB | NSB | $+0.6 \pm 0.13$ |
| 60295 | $\sim 17$ $\mu M^*$ | | | $-0.2 \pm 0.10$ |
| 60296 | $9.32 \pm 0.28$<br>nM | $56.9 \pm 1.6$ | $53.0 \pm 0.6$ | $+3.1 \pm 0.06$ |
| 60297 | $\sim 2$ nM* | | | $+5.1 \pm 0.21$ |

\* Complex profiles required fitting to a heterogeneous model; the tighter of the two $K_d$ values is shown.
NSB = non-specific binding to the reference sensor, precluding analysis
low = low binding signal, precluding analysis

As an orthogonal approach, we investigated binding affinity of the ten peptides by thermal shift assay (TSA). The melting temperature of PfUCHL3 is 51.6 °C and a range of melting temperature shifts was observed in the presence of each peptide **(Figure 2A)**. The largest stabilisation was seen upon addition of peptide 60297, which increased the melting temperature of PfUCHL3 by 5.1°C, followed by peptide 60296 which increased the melting temperature by 3.1 °C at 25 μM.

**Figure 2.**
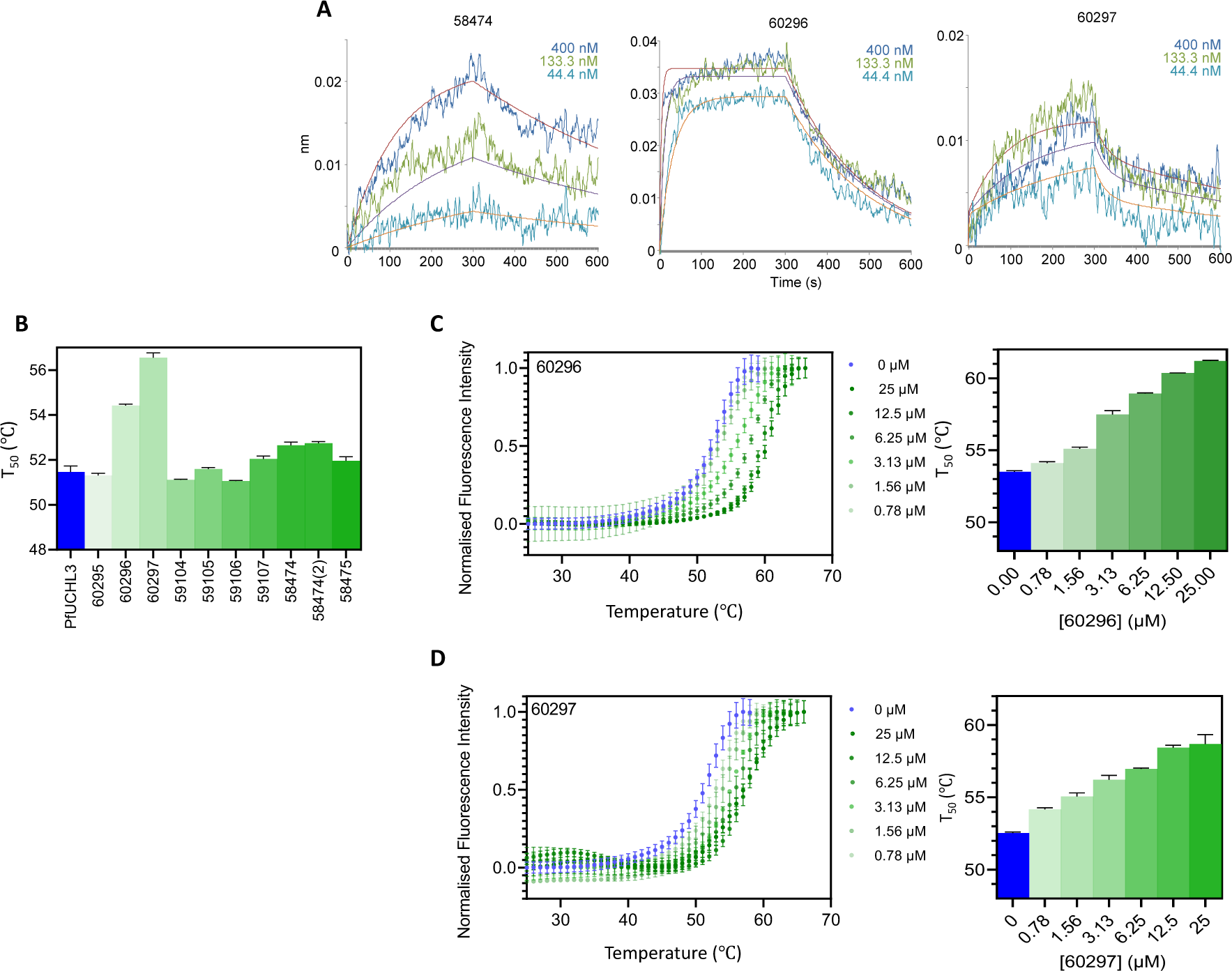
BLI and TSA analysis. A) BLI data: streptavidin-biotin immobilised protein dipped into peptides at the concentrations shown for 300s, then into buffer for 300s. Global fits are to a 1-site model for 58474 and 60296, and a 2-site model for 60297. B) Thermal shift data for 5 μM PfUCHL3 alone or incubated with each of the 10 peptides at 6 μM. C) Thermal shift data for PfUCHL3 (5 μM) alone (blue) incubated with serial 2-fold dilutions of peptide 60296 (C) and peptide 60297 (D) from 25 μM to 0.78 μM (dark green to light green).

Peptides 58474 and 58474(2), were the only other two peptides to cause a significant increase in the melting temperature (1.2 °C and 1.3 °C, respectively). Peptides 59107 and 58475 only increased the melting temperature by 0.5-0.6 °C. Peptides 59105 and 60295 had no significant effect, and 59104 and 59106 showed small decreases in the melting temperature of −0.4 °C. Thus, overall, the trend in melting temperatures rough follows that of the *K*_d_ values measured by BLI. To explore the effects of peptides 60296 and 60297 further, the dose dependence was measured. A concentration range of 48 pM to 25 μM was used with PfUCHL3 at a constant concentration of 5 μM, and the melting temperature was shown to increase in a dose-dependent manner **(Figure 2B,C)**.

### The peptides are potent and selective inhibitors of PfUCHL3-mediated ubiquitin hydrolysis

Having determined that the ten peptides bind to PfUCHL3, we next tested their ability to disrupt the enzyme’s deubiquitinating activity, as binding does not necessarily correspond to inhibition. Hydrolysis of the fluorogenic ubiquitin amido-methyl-coumarin (Ub-AMC) substrate (125 nM) by PfUCHL3 (31.25 pM) was assessed in the presence of each peptide at a concentration of 25 μM. Seven of the peptides showed no inhibition, two showed partial inhibition, and two showed complete inhibition. Ub-AMC assays were repeated for the four active peptides at different concentrations. In agreement with the BLI and TSA results, the two most potent inhibitors were peptides 60296 and 60297 which had IC50 values of 14.6 ± 2.4 nM and 1438 ± 371 nM respectively **(Figure 3)**. Importantly, all four peptides had no inhibitory effect on the human UCHL3 enzyme **(Figure 3E)**. Two control peptides were synthesised, peptides 62605 and 62606, in which the sequences were scrambled versions of peptides 60296 and 60297. Again, no inhibition was observed for these peptides even at 25 μM concentrations **(Supp Figure 2)**. Thus, the two lead peptides are sequence-and species-specific inhibitors of PfUCHL3. The selectivity for PfUCHL3 over HsUCHL3 is striking given the high sequence identity between the two proteins (**Supp Figure 3**) particularly in the active site and the ubiquitin substrate binding site; furthermore, the sequences of ubiquitin from the two species differs by only 1 amino acid, and thus the enzyme-substrate interfaces are very similar.

**Figure 3.**
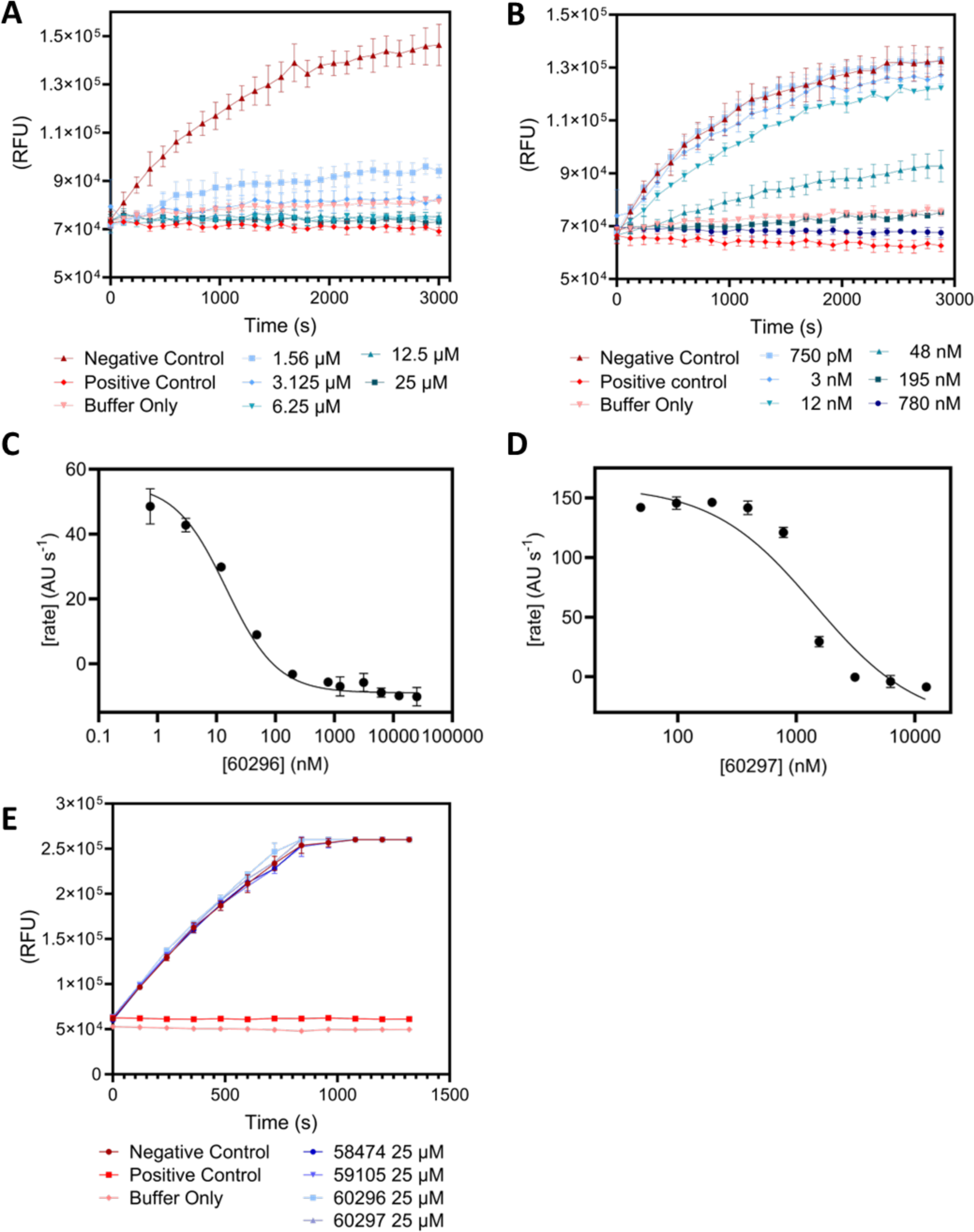
Ub-AMC assays. PfUCHL3 (31.25 pM) incubated with serial 2-fold dilutions of peptide 60296 (A) and 60297 (B). ‘Positive Control’ corresponds to N-ethylmaleimide (NEM) inhibited PfUCHL3; ‘Negative Control’ corresponds to PfUCHL3 only (without substrate, no peptide/inhibitor)’; ‘Buffer Only’ corresponds to the buffer system with no enzyme or peptides present. Plot of maximum rate versus peptide concentration for 60296 (C) and 60297 (D). Ub-AMC assay of the peptides (25 μM) incubated with HsUCHL3 (31.25 pM) to test species specificity (E).

### The peptides bind to the ubiquitin-interaction interface on PfUCHL3

To understand the mode of action underpinning the observed inhibition, we next performed NMR experiments. First, we assigned the 2D ^1^H-^15^N transverse relaxation optimised spectroscopy (TROSY) spectrum acquired with ^15^N-labelled PfUCHL3. We next incubated the protein with each of the two lead peptides, 60296 and 60297, at a 1:1 molar ratio. For both peptides a specific subset of amino acids displayed chemical shift changes with some displaying very large movements in their chemical shift values **(Figure 4)**. This result indicates that the peptide interaction is localised, potentially including a large number of aromatic side chains, and is not eliciting a global conformational change in the protein. The PfUCHL3 residues displaying a significant chemical shift upon addition of either peptide were mapped onto the ubiquitin-bound PfUCHL3 structure. This structure reveals two major interaction interfaces on PfUCHL3 for ubiquitin, a charged interface around the active site and a distant hydrophobic interface referred to as the ubiquitin-recognition site. The electrostatic active site interaction involves the insertion of the LRLRGG sequence at the C-terminus of ubiquitin into the active site via a tunnel created by the PfUCHL3 “crossover loop”. At the hydrophobic ubiquitin-recognition site, the sequence KTLTGK of ubiquitin forms a hairpin turn that inserts into a hydrophobic pocket on the surface of PfUCHL3. For binding of both peptides, we see that the PfUCHL3 residues that displayed the largest chemical shift changes map very closely to the hydrophobic ubiquitin-recognition site **(Figure 4)**.

**Figure 4.**
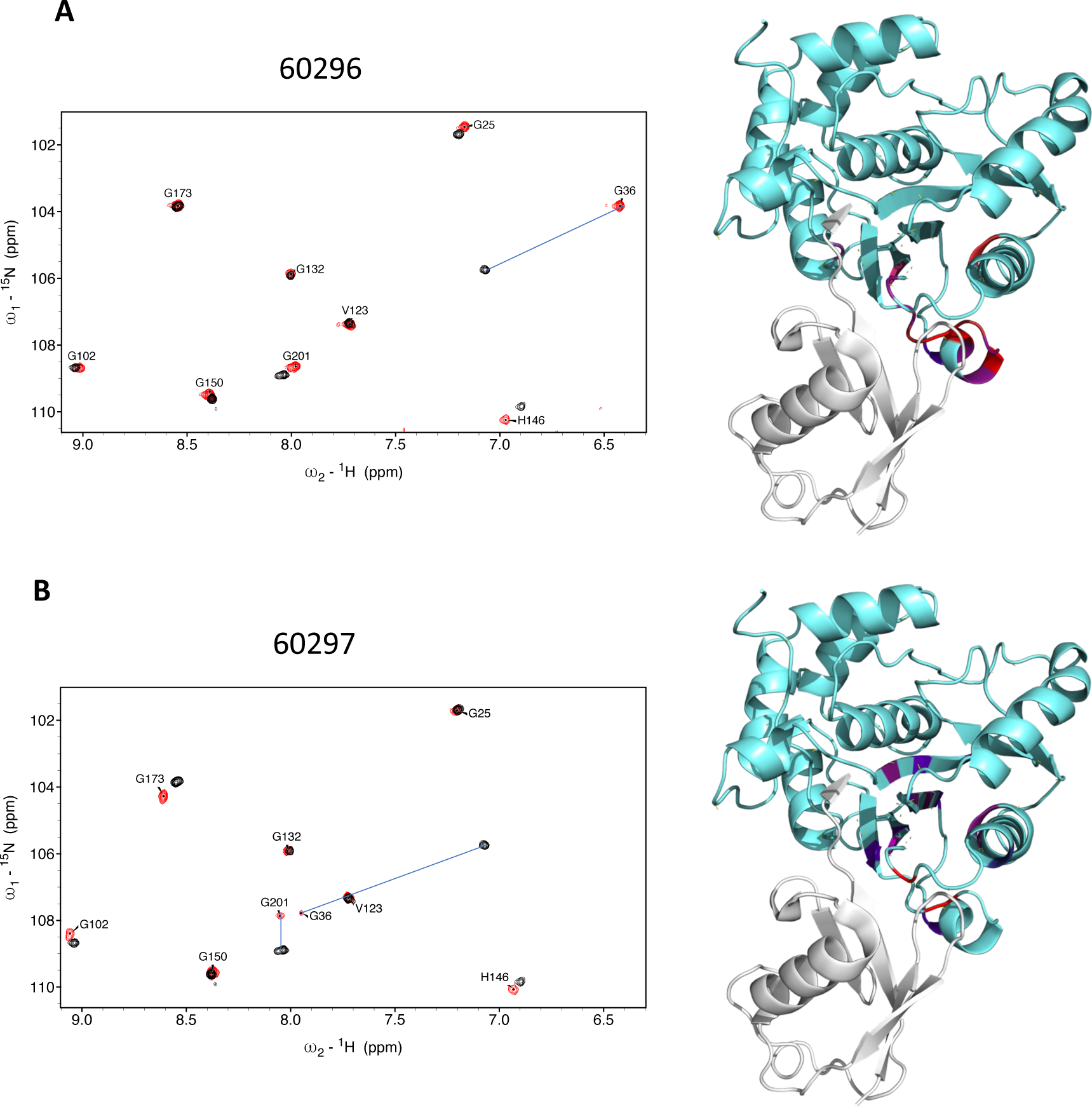
NMR spectroscopy of PfUCHL3 maps the binding site of the peptides. ^1^H-^15^N HSQC spectra of PfUCHL3 without (red) or with (blue) the addition of peptide 60296 (A) or peptide 60297 (B) (ratio 1:1). The changes in peak positions for residues that undergo changes in chemical shift of more than two standard deviations are indicated. Images on the right show the crystal structure of PfUCHL3 (cyan) in complex with Ub (grey) (PDB code: 2WDT [12]) with the PfUCHL3 residues that show significant chemical shift changes upon peptide binding highlighted in heat map coloration. See also Supp Figures 4 and 5.

In the case of peptide 60296, we observed extensive chemical shift changes in the ubiquitin-recognition site and adjacent β-sheet structure of PfUCHL3 upon peptide binding. It is possible that the peptide may be expanding the naturally occurring substrate-binding pocket to accommodate a larger binding interface between peptide 60296 and PfUCHL3. This slightly remodels the enzyme and induces structural changes in the β-sheet core, which is located in close proximity to the ubiquitin recognition region. The finding is consistent with the TSA data showing large thermal stabilisation of PfUCHL3 upon peptide binding and with 60296 being the most potent inhibitor.

Peptide 60297 induced similar changes in PfUCHL3, albeit to a slightly lesser extent. Thus, the peptides interact with the hydrophobic pocket and compete for binding with the KTLTGK sequence of the ubiquitin substrate, thereby inhibiting activity. Additional experiments were performed whereby peptide 60296 was incubated with PfUCHL3 at a 1:0.5 molar ratio. The results showed that the bound and unbound forms of the protein were in slow exchange, which is consistent with the slow off-rates observed in the BLI experiments.

### In silico modelling reveals that peptides block the interaction of Ub with PfUCHL3

To provide further atomic detail on the binding sites of the peptides, we combined the NMR results with computational modelling. Peptides 60296 and 60297 were initially modelled using published cyclic peptide structures (PDB code: 6u74 (34). Although the crystal structures of PfUCHL3 protein are available in apo (PDB code: 2WE6 (12) and holo (Ub-bound) (PDB code: 2WDT (12)) forms, both of them lack an ⍺-helical region (residues 59 to 76), which is 5 Å from the allosteric ‘ubiquitin-recognition site’. Hence, the AlphaFold (Uniprot ID: Q8IKM8) structure of PfUCHL3 protein was used to dock the peptides onto the binding site defined by the surface-exposed residues of PfUCHL3 with significant ^1^H chemical shift changes. The poses are similar for the two peptides **(Figure 5A),** and the interface overlaps with the ubiquitin substrate-binding interface, consistent with the NMR results. Both 60296 and 60297 structures have aromatic residues, W and Y, respectively, that anchor at the allosteric ‘ubiquitin-recognition site’ of the PfUCHL3 protein. The cyclisation of the peptides helps to lock them in a binding-competent conformation **(Figure 5B,C)**.

**Figure 5.**
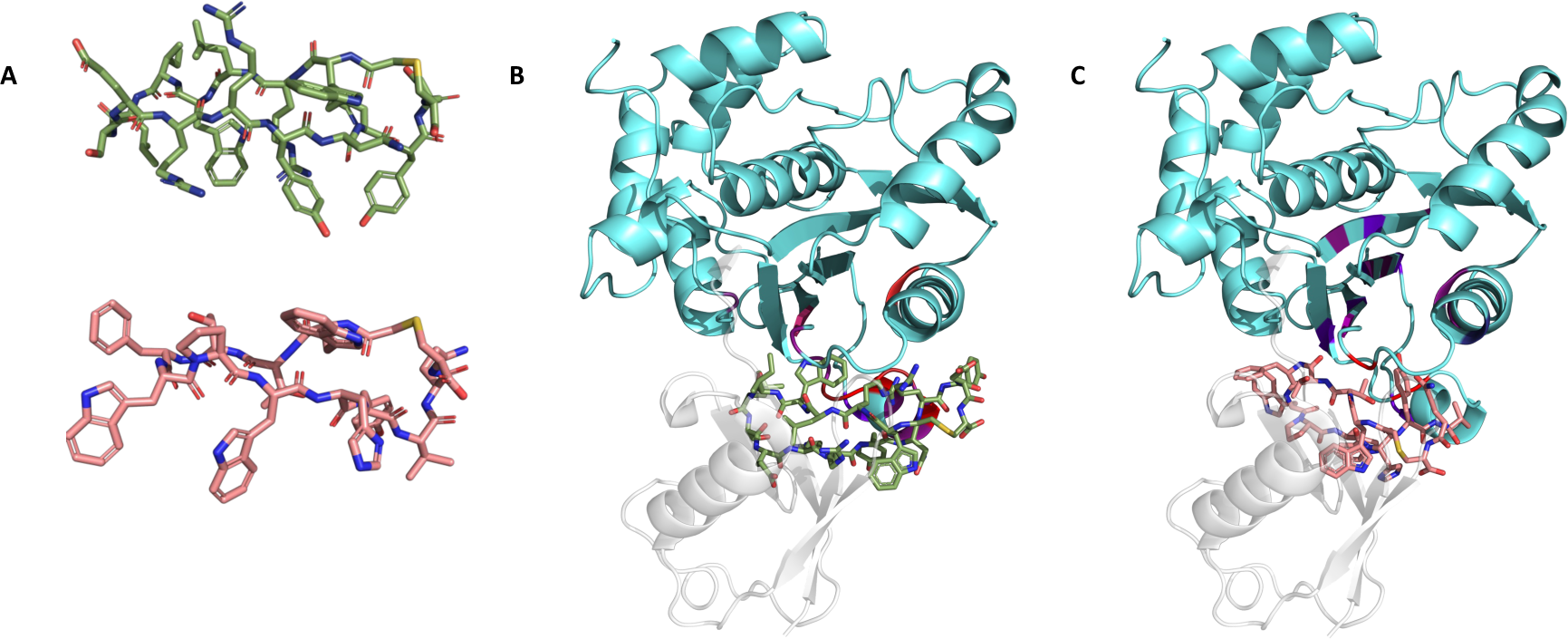
*In silico* modelling and docking. A) Cyclic peptides 60296 (top) and 60297 (bottom) modelled using Modeller v.9.25 program. PfUCHL3 (cyan) is shown in complex with the docked cyclic peptides 60296 (B) and 60297 (C) and with the Ub (PDB code: 2WDT, grey (12)). Amino acid residues of PfUCHL3 involved in the binding heatmap generated by NMR are highlighted.

## Discussion

As in all eukaryotes, the ubiquitin pathway is critical to *Plasmodium* parasites for their survival, replication, and cellular homeostasis. Undoubtedly, many of the pathway components have yet to be discovered owing to this parasite’s unique biology and lack of sequence homology to ubiquitin enzymes in higher eukaryotes. The majority of those that have been identified are essential for parasite viability, expressed across multiple life-cycle stages and possess only moderate identity to homologous human proteins. These points highlight the ubiquitin pathway as a promising target for the development of novel antimalarials.

In this study, we used the RaPID selection technique to identify cyclic peptide inhibitors of PfUCHL3, a dual activity DUB/deneddylating enzyme of *Plasmodium falciparum*, the deadliest malaria parasite. PfUCHL3 has been previously crystallized in both the apo and Ub-bound forms. The structures revealed two main points of contact between the enzyme and its cognate substrate: one in and around the active site and the other a hydrophobic pocket lined by Helix 7 and the loop preceding Helix 1, a signature interaction motif between UCH family proteases and Ub (12). Both human and *P. falciparum* UCHL3 structures reveal a cross-over loop that restricts access to the active site. Although it has been proposed that this loop may be flexible and able to swing aside to accommodate large, globular proteins [9], subsequent studies where the loop was cleaved or enlarged, significantly increased efficiency of di-Ubiquitin hydrolysis for both the parasite and human UCHL3 enzymes (11). These observations have led to the suggestion that the natural substrate for this enzyme is in fact small, with its main role being to replenish the free ubiquitin pool by cleaving adventitiously-generated ubiquitin adducts such as thiols or amines (35). Although the wider biological function of PfUCHL3 is still unknown, transcriptomic and proteomic analyses reveal its presence across multiple stages of *P. falciparum* development suggesting it is functional - and therefore targetable - throughout the parasite’s lifecycle (36–38). We identified two peptides that bind to PfUCHL3 with nanomolar affinity and are capable of inhibiting the deubiquitination activity of PfUCHL3 *in vitro* but not that of human UCHL3. NMR spectroscopy revealed that the peptides interact with the hydrophobic pocket proximal to the active site rather than the active site itself. Considering a peptide would likely have to be linear in order to access the catalytic region under the cross-over loop, it is perhaps unsurprising that a screen based on cyclic peptides with structural rigidity would select for other locations on the enzyme’s surface.

Modelling indicated that binding of the two lead peptides sterically blocks ubiquitin from adopting the necessary orientation to access the PfUCHL3 active site, thereby inhibiting the enzyme’s DUB activity. We are now working to assess peptide inhibition *in vivo*. Localisation of the human ortholog is predicted to be diffuse, spanning both cytoplasm and nucleus. Localisation studies for the PfUCHL3 are largely consistent with this distribution, but this would still require a peptide to cross three membranes to access the enzyme: the erythrocyte, the parasitophorous vacuole, and the parasite plasma membranes. Although this may potentially be challenging, cyclic peptides have been previously and successfully used as inhibitors of *P. falciparum* proteins *in vivo*. In particular, STAD-2, a hydrocarbon stapled peptide with antiplasmodial activity, was shown to access the cytoplasm of infected red blood cells (39). The exact mechanisms of how this peptide crosses the membrane has not been fully elucidated, but possible routes might include passive diffusion or uptake through endocytosis followed by endosomal escape. Indeed, there has been sustained interest in developing cell penetrating peptides capable of escaping from endosomes, particularly in the context of viral drug delivery (21). *Plasmodium* presents a slightly more complex situation in that endosomes are generated by the double invagination of the PV and plasma membranes and result in double-membraned structures. Drawing from insights amassed as part of these studies, next-generation PfUCHL3 inhibitors can be designed to incorporate additional unnatural amino acids and cell-penetrating tags to improve permeability. Moving forward, we aim to develop these inhibitors to disrupt parasite growth and as tools to elucidate the biological role of this enzyme in *Plasmodium* parasites.

## Materials and Methods

### Protein expression and purification

Immobilised metal affinity chromatography (IMAC) was used to purify proteins containing a His6-tag. 5 mL HisTrap excel columns (GE Healthcare, column pressure limit = 0.5 PSI) were used with flow rates at 6 ml/min and loaded either by hand or by using an AKTA Purifier (GE Healthcare) following manufacturers guidelines. Protein was loaded at 3 mL/min and eluted at 5 ml/min into 96 deep-well blocks. 20 mM imidazole was used for washing the matrix and 300 mM imidazole was used for eluting the bound protein. Eluted proteins had their purity checked by SDS PAGE and the purest fractions of POI were combined for further use.

For library screening, constructs with N-terminal 6xHis were cloned into expression vectors, extending the N-terminus with an Avi-tag (GLNDIFEAQKIEWHE), an amino acid sequence that can be specifically biotinylated by the enzyme BirA. PfUCHL3-His6-Avi plasmid was transformed into CD41 bacterial cells and induced with 1 mM IPTG overnight. Cells were harvested, resuspended in lysis buffer (50 mM Tris-HCl buffer pH 8.0, 150 mM NaCl, 15 mM imidazole, 2 mM EDTA, and 1 mM TCEP) and lysed using an EmulsiFlex C5 homogenizer (Avestin) at 10-15,000 pounds per square inch followed by passage over a HisTrap column (GE Healthcare) and elution with 300 mM imidazole in lysis buffer. Peak fractions were collected and diluted to 75 mM NaCl using IEX buffer (50 mM Tris-HCl buffer pH 8.0, 2 mM EDTA, and 1 mM TCEP), followed by further purification with a MonoQ 10/100 GL column (GE Healthcare) using a 0.75–1 M salt gradient in the same buffer. Fractions containing PfUCHL3 were then pooled, concentrated, and injected onto a HiLoad 16/600 Superdex 75 pg column (GE Healthcare) equilibrated with storage buffer (25 mM Tris-HCl buffer pH 8.0, 200 mM NaCl, 2 mM EDTA, and 1 mM DTT). Peak fractions were pooled, concentrated, and flash-frozen in aliquots. After purification, recombinant BirA was added to the purified complexes in the presence of biotin to produce the biotinylated complex, which was suitable for immobilization and mRNA display screening.

For PfUCHL3, Biomix A (0.5 M bicine buffer, pH 8.3), Biomix B (100 mM ATP, 100 mM Mg(OAc)2 and 500 μM D-biotin), concentrated defrosted PfUCHL3-AviTag (375 μM), PfUCHL3 buffer (25 mM Tris-HCl buffer, 200 mM NaCl, 1 mM DTT, 2 mM EDTA, pH 8.0) and BirA (50 μM) were mixed in a ratio of 2.5: 2.5: 2.5: 15: 1.. The reaction was incubated on tube rollers (R.T.P, 18 hours) and loaded onto a HiLoadTM 16/600 Superdex 75 pg column pre-equilibrated with PfUCHL3 buffer. Protein purity was checked by 10% SDS-PAGE. Biotinylation was checked by Western blot, and protein was concentrated, and flash frozen.

All biochemical and biophysical assays used UCHL3 proteins lacking the Avi tag, which were purified essentially as described for their Avi-tagged counterparts, with the exception that the 6xHis tag was cleaved from PfUCHL3 by addition of TEV protease following HisTrap purification and repassage of the product over the HisTrap column to remove the cleaved tag and protease. Peak fractions were pooled, concentrated, and flash-frozen in aliquots.

### Screening of PfUCHL3 binding macrocyclic peptides with the RaPID system

*In vitro* selections of PfUCHL3-binding macrocyclic peptides using the RaPID system was performed as previously reported (40) with slight modification. Briefly, the initial random mRNA library was transcribed and ligated to a puromycin linker primer via T4 ligase for 30 min at 25 °C and extracted with phenol/chloroform and ethanol precipitated. A 150 µL translation reaction using the methionine-deficient FIT system (41) and a 50 µM concentration of ClAc-D-Trp-tRNA^fMet^_CAU_ were used to convert the mRNA library into a library of peptide-mRNA fusions. The translation was performed at 37 °C for 30 min followed by a 25 °C step for 12 min to enhance the formation of the peptide-mRNA fusions. 30 μl of 100 mM EDTA was added to dissociate ribosomes and the peptide-mRNA fusions were incubated at 37 °C for 30 min to allow the thioether cyclization to approach completion.

The fused peptide–mRNA was subsequently reverse transcribed using MMLV RT Rnase H-(Promega) for 1 h at 42 °C. The resulting peptide-mRNA-DNA fusions were then incubated with PfUCLH3 immobilized on Dynabeads M-280 streptavidin (Invitrogen) for 30 min at 4 °C. The resultant complementary DNAs were eluted by mixing with PCR reaction buffer and heating at 95°C for 5 min, followed by immediate separation of the supernatant from the beads. A small fraction of the cDNA and input were allocated to real-time PCR quantification using a LightCycler 2.0 (Roche); the remainder was amplified by PCR. The resulting duplex DNAs were purified by phenol–chloroform extraction and ethanol precipitation and transcribed into mRNAs for the next round of selection. From the second round of selection, the translation was performed at 5 µL scale, and six times of pre-clear steps were added as negative selection preceding the positive selection steps using 1 µL each of untreated and biotin bound Dynabeads M280. Finally, the observed DNA populations appearing at fifth, sixth, and seventh rounds were subjected to next generation sequencing using the MiSeq sequencing system (Illumina).

### Chemical synthesis of peptides

Macrocyclic peptides were synthesized and purified by Cambridge Peptides and Peptide Synthetics.

### Ub-AMC deubiquitination (DUB) assay

DUB assays were performed in triplicate in 384-well plates (Corning) using a CLARIOstar plus plate reader. A stock mix of either PfUCHL3 or HsUCHL3 was prepared in the presence of either DMSO (mock) or cyclic peptide. Before making the stock mix, the enzyme complex and peptide were diluted in reaction buffer (50 mM Tris-HCl buffer, pH 7.5, 150 mM NaCl, 1 mg/ml BSA, 0.1% DMSO, and 1 mM TCEP) to 4x the desired final peptide/protein concentration (so as to keep DMSO concentration constant in all reactions). Protein and peptide were mixed 1:1 and incubated for 10 min at room temperature, with each enzyme complex at 2x the desired final concentration. We added 10 μL of the enzyme:peptide mixture to 10 μL of 250 nM Ub-AMC (7-amino-4-methylcoumarin) (Boston Biochem). DUB activity was measured by monitoring the increase in AMC fluorescence in a plate reader. The kinetic data at each concentration of peptide was corrected for background and then the initial rate of reaction was determined. The rate of reaction was plotted against concentration of peptide to determine the IC50 using the following equation:

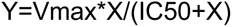

Where Vmax is the reaction rate in the absence of inhibitor, X is the concentration of the peptide inhibitor, and IC50 is the peptide concentration when the rate was inhibited by the peptide by 50%. The data were fitted using Prism (Graphpad).

### Biolayer interferometry (BLI)

BLI experiments were performed using a ForteBio Octet Red96 instrument at 25 °C. Measurements were performed in black 96-well plates with a well volume of 200 μl. Biotinylated Avi-tagged PfUCHL3 was prepared in a buffer containing 50 mM Tris-HCl buffer, 200 mM NaCl, 0.5 mM TCEP, pH 7.5, and loaded at 1 μg/mL for 60s. Peptides were diluted to a 4 μM stock concentration in the same DMSO:water mix in which they were resuspended from their stock powder (approximately 4%). The final dilutions (to 400 nM, 133.3 nM and 44.4 nM) were performed in the same buffer as for the PfUCHL3. Streptavidin-coated tips were used with a new tip for each measurement, because regeneration with PfUCHL3 attached to the tip was not possible. The association and dissociation times were 300s. All data analysis was performed with the instrument software.

### Thermal shift assay

Thermal shift assays were performed using a Bio-Rad CFX Connect qPCR instrument, whereby the unfolding is detected by fluorescence of the hydrophobic dye Sypro Orange that binds to the unfolded state of the protein. The experiments were performed in clear-bottom, half-volume, 96-well plates using final well volumes of 25 μL, PfUCHL3 concentration of 5 μM, peptide concentrations ranging from 47 pM to 25 μM, and Sypro Orange:protein concentration ratio of 1:400. PfUCHL3, Sypro orange and peptide were diluted to 5x concentration in buffer, 1:2000 concentration in buffer, and 4% DMSO respectively, before being mixed in the well to a final 1x concentration in buffer. For the PfUCHL3-only control, the peptide was replaced with 4% DMSO and for the buffer control, the PfUCHL3 was replaced with buffer in the sample mix. The samples were incubated at 20 °C for 2 minutes before increasing by 0.5 °C every 30 seconds up to 90 °C. At each temperature the fluorescence intensity was measured using an excitation wavelength of 471 nm and an emission wavelength of 570 nm. Data analysis was performed using the GraphPad Prism software.

### NMR spectroscopy

^15^N-labelled and ^13^C/^15^N-labelled PfUCHL3 proteins were produced by growing cells in K-MOPS minimal media supplemented with ^15^N-labelled ammonium chloride and ^13^C-glucose to obtain the desired isotope-labelled protein. The proteins were purified as described above. NMR spectra were recorded at the MRC Biomolecular NMR Centre at the Crick Institute (London) on a 950 MHz (Bruker Avance III HD 950) and a 700 MHz (Bruker Avance III HD 700) equipped with a triple-resonance cryoprobes on a 90% H2O, 10% D2O 100 μM sample containing 20 mM Tris-HCl buffer pH 7.5, 5 mM NaCl, 0.5 mM EDTA, 0.5 mM TCEP at 25 °C. Backbone assignments were carried out using HNCO, HN(CA)CO, HNCA, HN(CO)CA, HNCACB and CBCA(CO)HN 3D heteronuclear NMR experiments on ^13^C/^15^N-labelled samples using standard Bruker pulse programs. Topspin (Bruker) was used for data processing and the POKY software package (42) was used for data analysis. Backbone assignments were initially made automatically using MARS (43) and completed manually. Protein:peptide samples were prepared in a 1:1 ratio unless stated otherwise. Chemical shift perturbations (CSPs) induced by peptide binding were calculated as the root mean square deviation (RMSD) of the changes of the H and N chemical shifts.

### In silico modelling

To build models of our cyclic peptides, we searched the PDB for structures of cyclic peptides with disulphide linkages similar to those used here. The cyclic peptide in the highest resolution (1.85 Å) complex was one in complex with acetyllysine-binding bromodomain proteins BRD4 (PDB ID: 6u74 (34)) and was used as a template to build all three cyclic peptides of interest (59105, 60296, and 60297) using Modeller v.9.25 program (44). The model with the lowest discrete optimised protein energy (DOPE) score (45) was selected. Next, a control docking experiment was performed to validate the docking protocols in Autodock Vina (46) using the BRD4-cyclic peptide complex (PDB ID: 6u74). The centroid of the grid box sized 35 Å x 35 Å x 35 Å was defined by the BRD4 residues within 4 Å of the cyclic peptide. RMSD of the atoms that make important interactions, namely O and N atoms, between the docking pose and the cyclic peptide from the crystal structure was calculated. The docking poses from Autodock Vina were then reranking using the nnscore (47) and rfscore (48) in the ODDT package (49). The scoring function that correlated with the RMSD was chosen. The docking between the cyclic peptide and BRD4 correctly modeled the binding interaction to within 0.4 Å of that in the published crystal structure.

The docking was then performed on the AlphaFold (50) model structure of PfUCHL3 protein (Uniprot ID: Q8IKM8) and the cyclic peptides. The same grid box size but with the centroid defined by residues that marked as 1 for LW or higher than 0.15 for dH in the NMR experiment was used.

## Acknowledgments

We thank members of the Artavanis-Tsakonas and Itzhaki labs for their help throughout this project and for useful discussions. Particular thanks go to Dr Maria Zacharopoulou for help with manuscript preparation. This work was supported by an MRC DTP studentship awarded to HRK and a Biotechnology and Biological Sciences Research Council (BBSRC) project grant awarded to KAT (BB/R001642/1) and the Japan Society for the Promotion of Science (JSPS) Grant-in-Aid for Specially Promoted Research (JP20H05618) to HS.

---

**Supplementary Figure 1.**
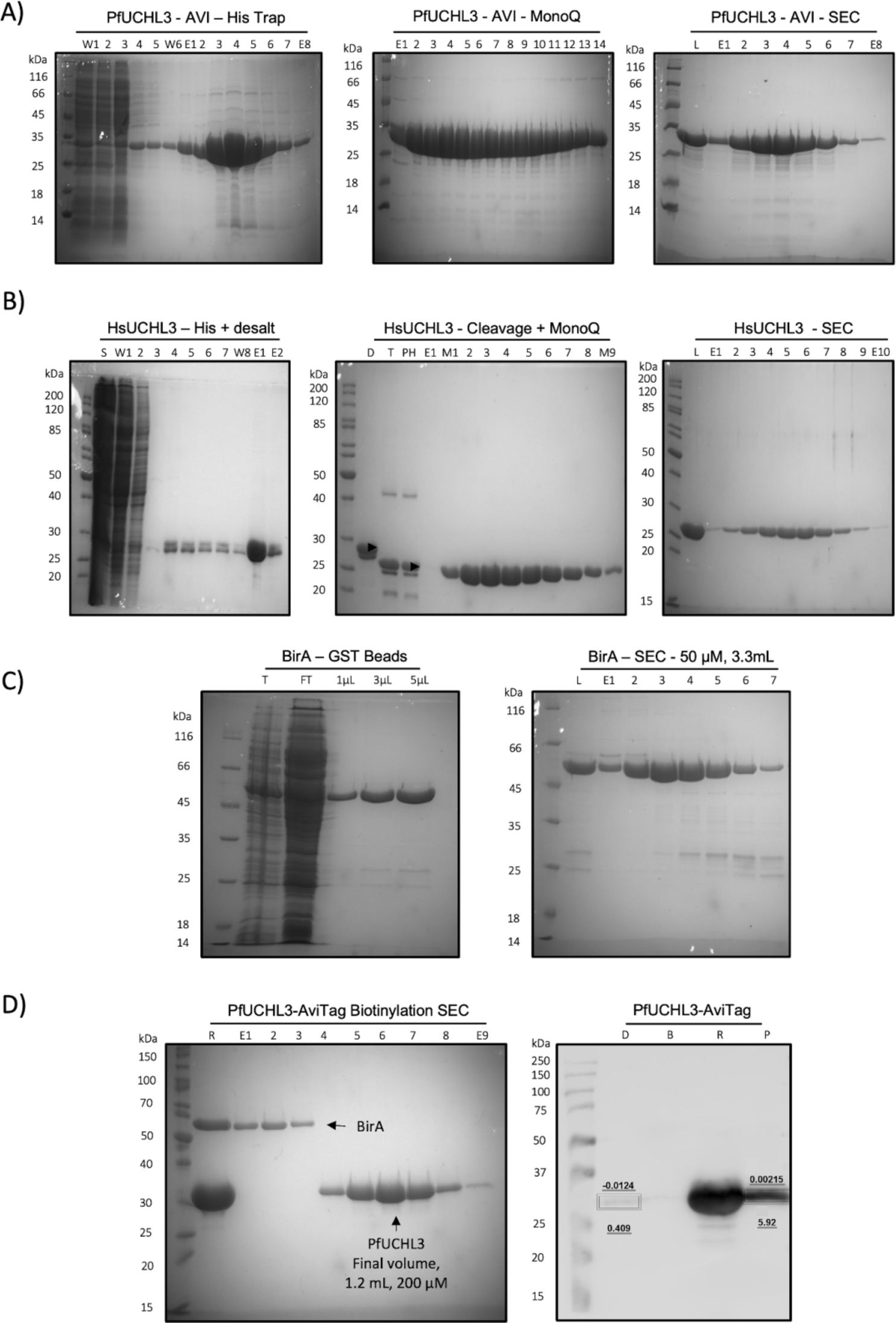
A) SDS PAGE gels showing the purification of PfUCHL3-AviTag. Left: His-trap purification; middle: ion-exchange step; Right: size-exclusion chromatography. B) The same gels as in (A) for the purification of HsUCHL3-AviTag. C) SDS PAGE gels showing the purification of BirA-GST tag. Left: purification by GST beads; Right: size-exclusion chromatography. D) Left: SDS PAGE of the purification of the PfUCHL3 enzyme from the biotinylation mix; Right: Western blot showing the successful biotinylation of PfUCHL3 and capture of the protein on streptavadin beads.

**Supplementary Figure 2.**
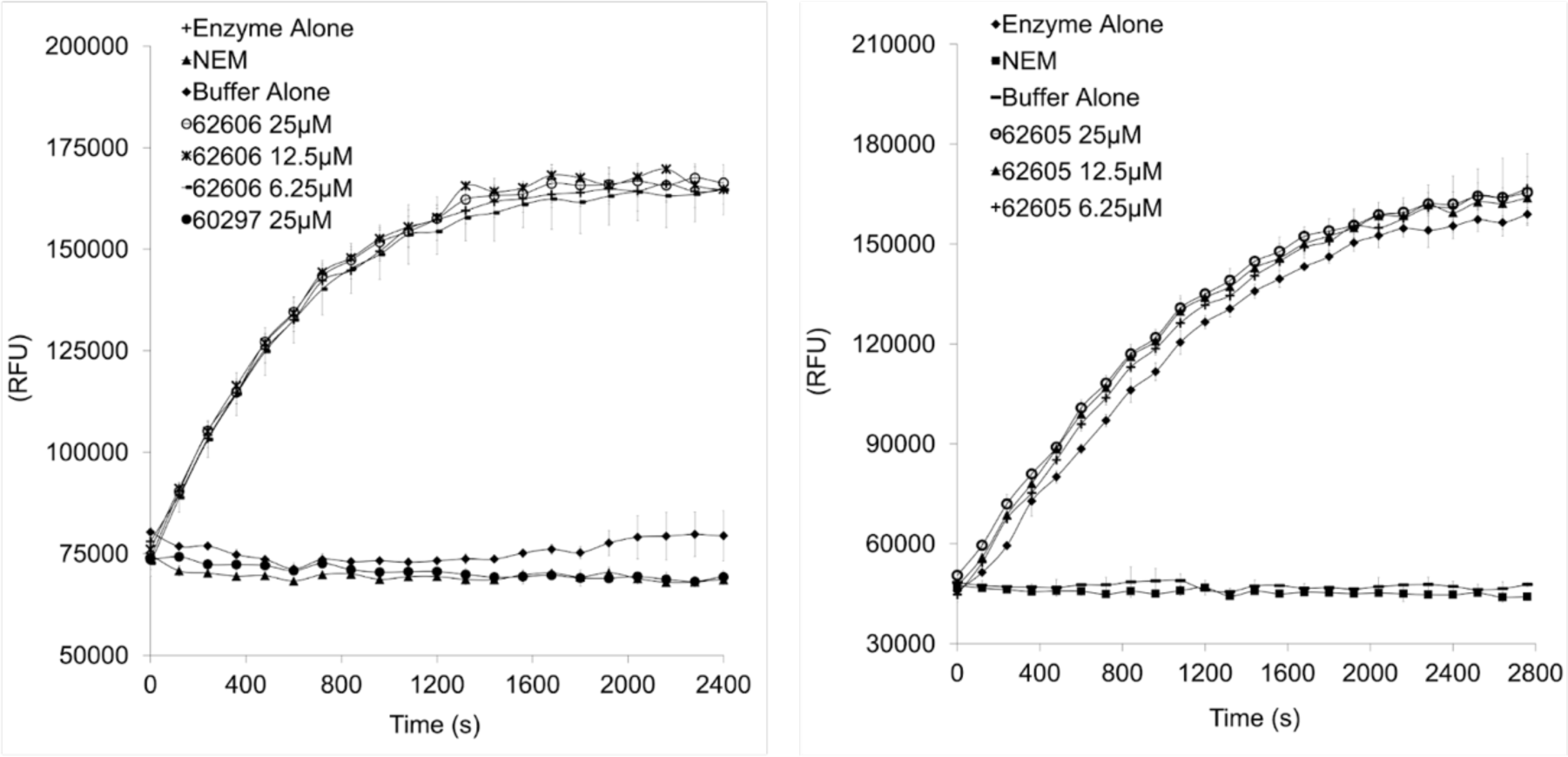
Ub-AMC assays of PfUCHL3 (31.25 pM) incubated with the scrambled peptides 62606 and 62605 (25 μM).

**Supplementary Figure 3.**
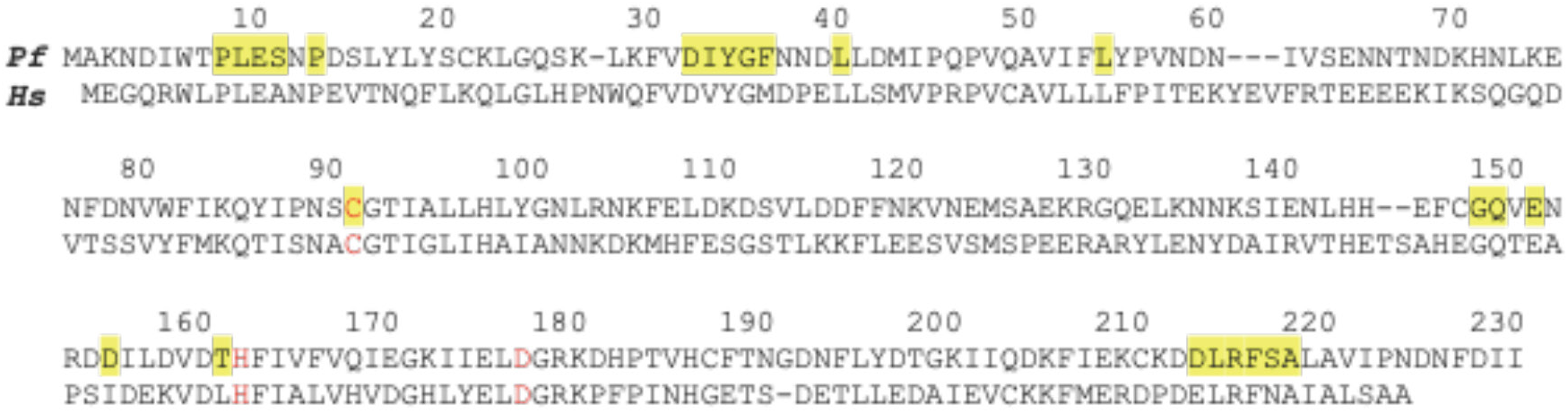
Sequence alignment of PfUCHL3 and HsUCHL3. Catalytic site residues are in red. Residues in the ubiquitin-binding site are highlighted in yellow.

**Supplementary Figure 4.**
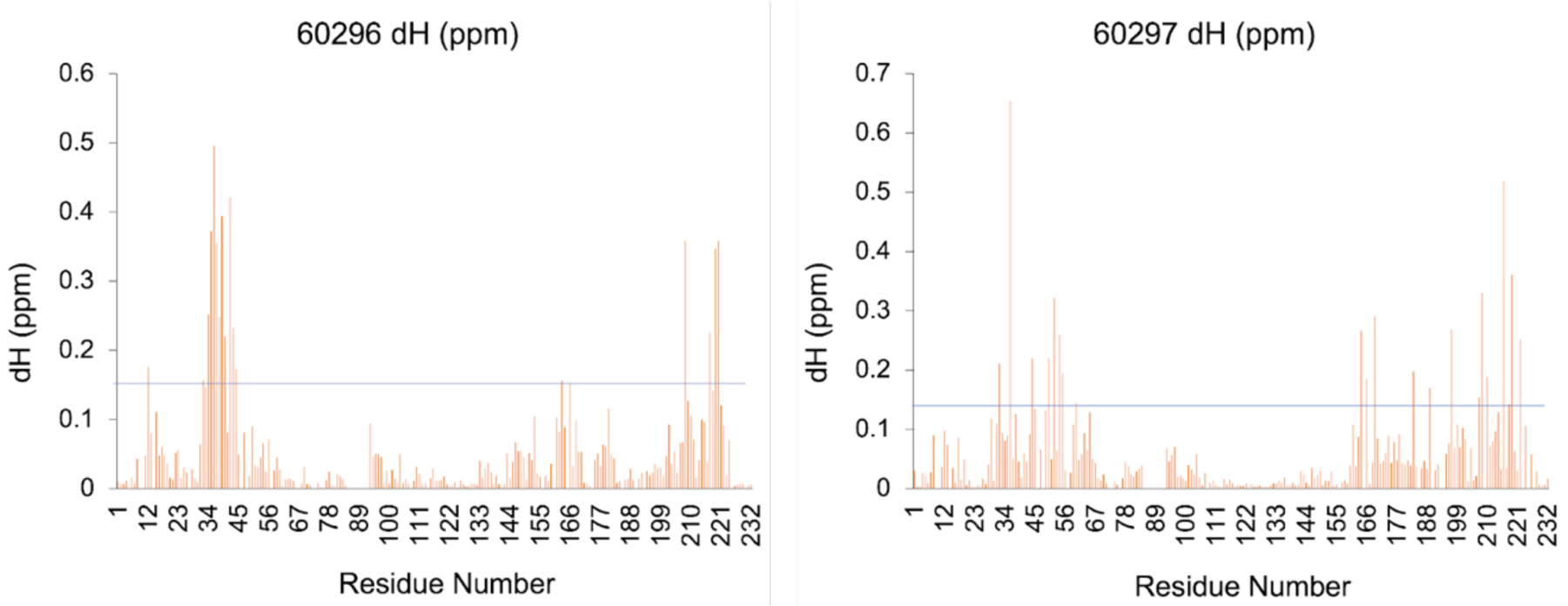
Diagram showing the changes in chemical shifts of the amide protons induced in the spectrum of ^15^N-labelled PfUCHL3 upon binding of the Peptide 60926 (left) and Peptide 60297 (right). The horizontal line indicates twice the calculated standard deviation in chemical shifts.

**Supplementary Figure 5.**
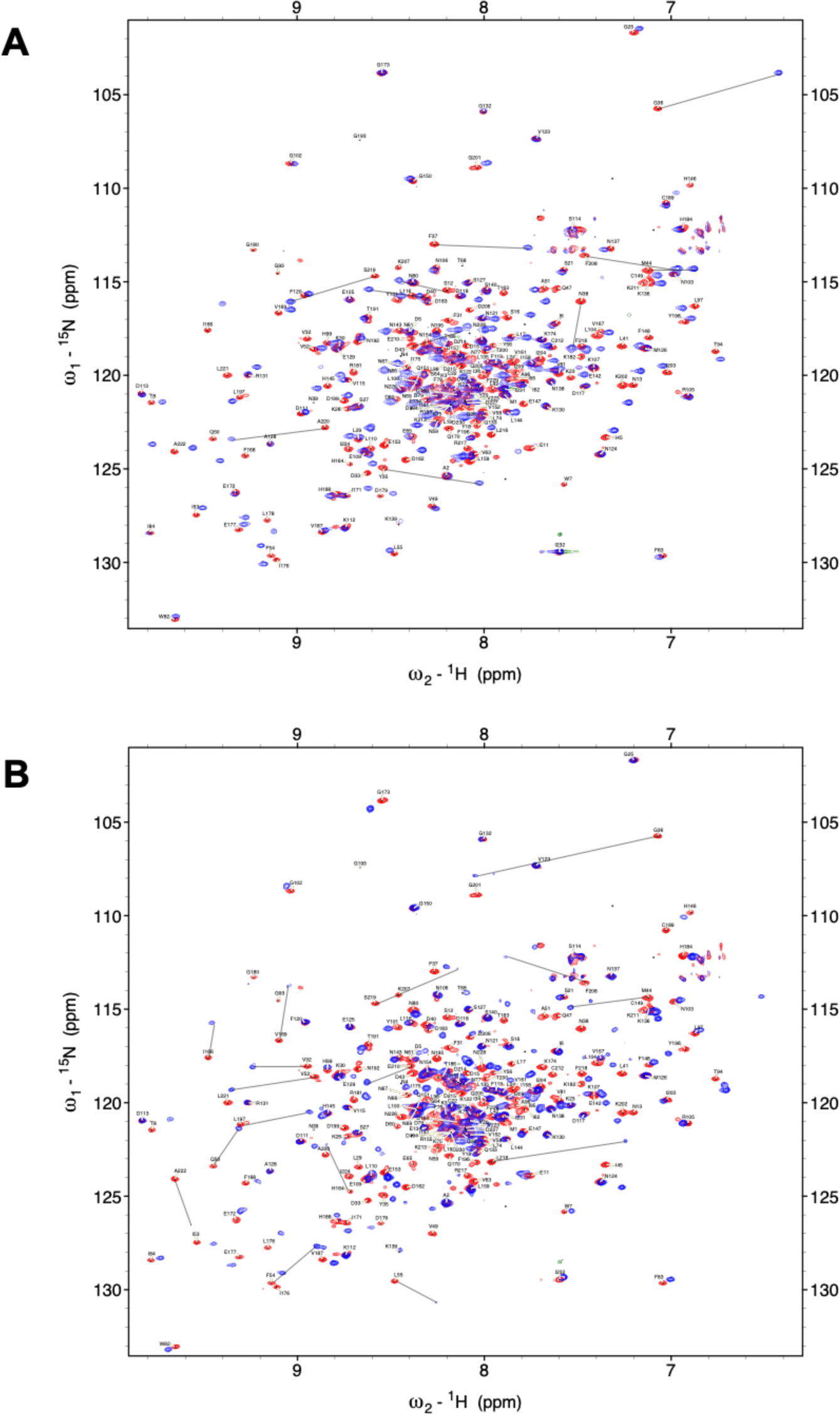
NMR spectroscopy of PfUCHL3 maps the binding site of the peptides expanded view. ^1^H-^15^N HSQC spectra of PfUCHL3 without (red) or with (blue) the addition of peptide 60296 (A) or peptide 60297 (B) (ratio 1:1). The changes in peak positions for residues that undergo changes in chemical shift of more than two standard deviations are indicated. Images on the right show the crystal structure of PfUCHL3 (cyan) in complex with Ub (grey) (PDB code: 2WDT [12]) with the PfUCHL3 residues that show significant chemical shift changes upon peptide binding highlighted in heat map coloration.

